# Lazarus effects: the frequency and genetic causes of *Escherichia coli* population recovery under lethal heat stress

**DOI:** 10.1101/058479

**Authors:** Shaun M. Hug, Brandon S. Gaut

**Affiliations:** Department of Ecology and Evolutionary Biology, U.C. Irvine, Irvine, CA

**Keywords:** Lazarus effect, heat stress, antagonistic pleiotropy, standing variation, population recovery

## Abstract

Sometimes populations crash and yet recover before being lost completely. Such recoveries have been observed incidentally in evolution experiments using *Escherichia coli*, and this phenomenon has been termed the “Lazarus effect.” To investigate how often recovery occurs and the genetic changes that drive it, we evolved ~300 populations of *E. coli* at lethally high temperatures (43.0°) for five days and sequenced the genomes of recovered populations. Our results revealed that the Lazarus effect is uncommon, but frequent enough, at ~9% of populations, to be a potent source of evolutionary innovation. Population sequencing uncovered a set of mutations adaptive to lethal 43.0°C that were mostly distinct from those that were beneficial at a high but nonlethal temperature (42.2°). Mutations within two operons—the heat shock *hslUV* operon and the RNA polymerase *rpoBC* operon—drove adaptation to lethal temperature. Mutations in *hslUV* exhibited little antagonistic pleiotropy at 37.0°C and may have arisen neutrally prior to subjection to lethal temperature. In contrast, *rpoBC* mutations provided greater fitness benefits than *hslUV* mutants, but were less prevalent and caused stronger fitness tradeoffs at lower temperatures. Recovered populations fixed mutations in only one operon or the other, but not both, indicating that epistatic interactions between beneficial mutations were important even at the earliest stages of adaptation.

## INTRODUCTION

The fossil record contains numerous examples of organisms that appear to have gone extinct for long stretches of geologic time, only to reappear in the fossil record later. One common explanation for these so-called “Lazarus” taxa is that the fossil record is incomplete, and fossil data are missing for lineages that always existed (Jablonski, 1986). It has also been suggested that the incompleteness of the fossil record for Lazarus taxa is not due to the loss of fossils or their inability to form, but rather extremely low population sizes. That is, Lazarus taxa do not suddenly reemerge in the fossil record due to missing data, but because their populations were actually on the verge of extinction, only to rebound later (Wignall and Benton, 1999).

A similar phenomenon has been observed in laboratory evolution experiments using bacteria, and it has been termed the Lazarus effect (Bennett and Lenski, 1993; Mongold *et al.*, 1999; Rodríguez-Verdugo *et al.*, 2014). When grown under lethal temperatures, bacterial populations decline in size over time, often to levels that are nearly or completely unquantifiable. However, some among the remaining survivors may acquire beneficial mutations that rescue the population and enable survival. These individuals reproduce, spreading the beneficial mutations through the population and restoring the population to sizes of similar magnitude to those observed before near-extinction.

Following the chance evolution of such a Lazarus population in an evolution experiment (Bennett and Lenski, 1993), Mongold et al. (1999) characterized patterns of evolutionary recovery at a lethal temperature of 44° using ancestral strains of *Escherichia coli* B that had adapted to low-glucose medium at 32°, 37°, and 41-42° (Mongold *et al.*, 1999). They found that Lazarus events at 44° occurred only in populations derived from ancestors already adapted to high temperature (41-42°), suggesting that pre-adaptation aids adaptive recovery under lethal temperature conditions. Moreover, they found that Lazarus mutants exhibited a fitness cost at elevated, but nonlethal temperatures, supporting the idea that some Lazarus mutations confer a fitness benefit in one environment, but a fitness cost in another: a phenomenon known as antagonistic pleiotropy (Williams, 1957; Cooper and Lenski, 2000; MacLean *et al.*, 2004).

Here we revisit the Lazarus effect, in part to determine whether Lazarus mutations are similar to, or distinct from, adaptive mutations that accumulate under thermal stress. For the comparison to thermal stress, we rely on an experiment by Tenaillon et al. (2012) that uncovered the genetic basis of adaptation to a high but sustainable temperature (42.2°). The experiment of Tenaillon et al. (2012) began with a strain of *E. coli* B that was adapted to low-glucose medium and 37.0° (*E. coli* B strain REL1206) (Tenaillon et al., 2012). The ancestral REL1206 strain was inoculated into 115 independent populations that were then evolved for 2,000 generations at 42.2°. Genome sequencing of a single clone from each population revealed >1,000 putatively adaptive mutations that fell into two major genetic pathways to high-temperature adaptation. The two pathways were exemplified by mutations in either *rpoB*, which encodes a subunit of RNA polymerase, or *rho*, which encodes a major transcriptional termination factor.

Subsequent work has focused on some of the *rpoB* mutations that occurred during adaptation to thermal stress. Rodríguez-Verdugo et al. (2014) engineered single *rpoB*mutations into the REL1206 background and found that they imparted an average fitness gain of ~22% at 42.2°. However, these same mutations tended to exhibit a fitness tradeoff at lower temperatures (<20.0°), consistent with the effects of antagonistic pleiotropy (Rodríguez-Verdugo *et al.*, 2014). It was during the course of this experiment, while investigating the fitness of clones at high temperature, that Rodríguez-Verdugo et al. (2014) observed Lazarus events at otherwise lethal temperatures in ancestral lines, despite the fact that the ancestor had not been pre-adapted to extreme temperature conditions.

In this study, we perform *E. coli* growth experiments to better understand the dynamics, genetics, and fitness consequences of Lazarus events. Beginning with an ancestor derived from a single-colony of *E. coli* B strain REL1206, we carry out replicated evolution experiments at two temperatures (43.0° and 44.0°) that typically result in population extinction under our growth conditions. After observing and noting the frequency of Lazarus events, we save populations that have rebounded from the brink of extinction, sequence their genomes from population samples, and determine the fitness of the Lazarus populations relative to the ancestor. Armed with genetic and fitness data, we seek to address three questions. The first is whether population recovery involves a distinct set of mutations relative to experiments at high but sustainable temperatures, as suggested by the fitness dynamics of Mongold et al. (1999). The second concerns the likely genetic drivers of adaptation, as opposed to hitchhikers. The discrimination of drivers and hitchhikers is a major challenge facing experimental evolution studies (Rosenzweig and Sherlock, 2014), but the short-term nature of our experiment permits novel insights into this distinction. The third question reflects the fitness effects of Lazarus populations and whether they exhibit tradeoffs at elevated, but nonlethal temperature (42.2°) or at the ancestor’s optimal temperature of 37.0°. Finally, combining genetic and fitness data allow us to comment on three important phenomena: antagonistic pleiotropy of adaptive mutations, the role of standing variation in population rescue, and the potential mechanism of population recovery.

## MATERIALS AND METHODS

**Lazarus Ancestral Stock:** A frozen glycerol stock was prepared from a single colony of *Escherichia coli* B strain REL1206 possessing a neutral Ara-marker. This strain had been propagated previously at 37.0 ° for 2,000 generations in Davis minimal medium supplemented with glucose at 25 mg/L (DM25), and was thus well adapted to the growth medium (Lenski *et al.*, 1991). To isolate the single colony, REL1206 was streaked from frozen onto a tetrazolium-arabinose (TA) plate and incubated overnight at 37.0°. The single colony was inoculated into Luria-Bertani medium (LB) and grown overnight. To prepare a frozen reference stock, 900 μL of culture was mixed with 900 μL of 80% glycerol and frozen at −80°. We term this REL1206 frozen stock the “Lazarus ancestor” (Figure 1). A backup Lazarus ancestor stock was prepared from the same LB culture.

**Figure 1:**
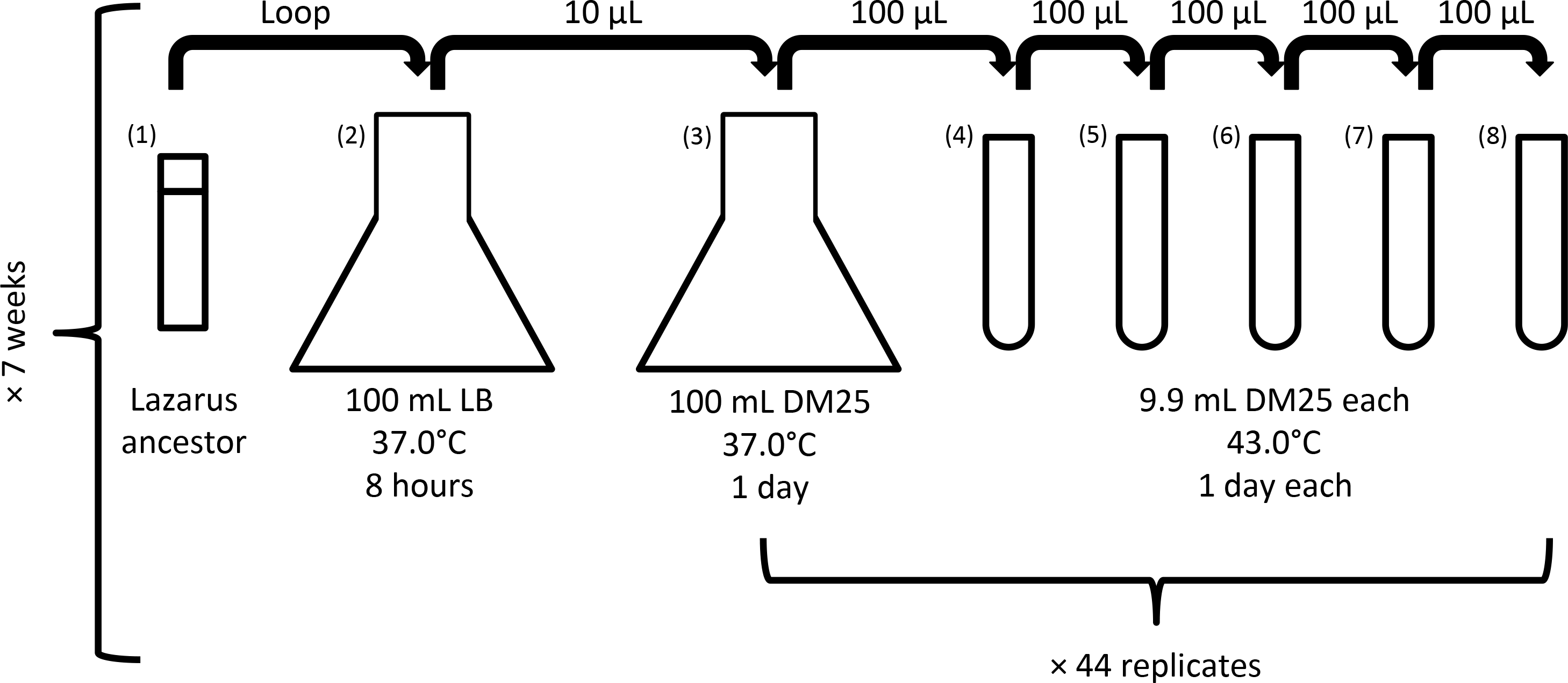
Experimental design for producing and observing Lazarus events. Bacteria were propagated from frozen through two flasks to acclimate them and to produce enough cells for experimental replication. Samples of flask culture were then propagated through culture tubes in 44 replicates for a total of five days. This procedure was repeated across seven different weeks.

**Lazarus Growth Experiments:** As is common practice (Bennett and Lenski, 1993; Lenski and Travisano, 1994; Rodriíguez-Verdugo *et al.*, 2014; Hug and Gaut, 2015), we first acclimated the Lazarus ancestor (REL1206) to mild laboratory conditions to allow it to recover from being frozen. The Lazarus ancestral stock was inoculated into 100 mL LB and grown for eight hours in an Infors HT Minitron incubator at 37.0° and 120 RPM (Figure 1). 10 μL of this culture was then inoculated into 100 mL DM25 and grown for 24 hours in an Infors HT Minitron incubator at 37.0° and 120 RPM. We inoculated 100 μL of the 37.0°, DM25 culture into each of 44 culture tubes containing 9.9 mL DM25. An additional four culture tubes containing 9.9 mL DM25 were used as contamination controls and cell density blanks. The total set of 48 tubes were placed into an Innova 3100 water bath shaker (New Brunswick Scientific) and grown for 24 hours at 120 RPM and at the experimental temperature of 43.0° or 44.0°. Tube cultures were serially propagated over the course of five days by inoculating 100 μL of culture into 9.9 mL DM25 after each day of growth (Figure 1).

On each of the five days, 50 μL of each culture (including blanks) was also inoculated into cuvettes containing 9.9 μL Isoton II Diluent (Beckman Coulter). These samples were analyzed using a Multisizer 3 Coulter counter (Beckman Coulter) to determine cell densities (particles/mL). The average particle density of the four blanks was subtracted from each sample’s cell density. We defined a Lazarus event as a population whose cell density increased by at least an order of magnitude over the previous day’s measurements.

Following cell density measurement, Lazarus populations were saved as frozen glycerol stocks on their initial day of recovery, as well as any subsequent days. To prepare frozen stocks of Lazarus recovery events, 900 μL of culture was mixed with 900 μL of 80% glycerol and frozen at −80°.

**DNA Extraction and Sequencing:** Most samples for DNA extraction and sequencing were derived from day five of the experiment; two samples (#1 and #19) were derived from day four. Each of the frozen Lazarus samples was inoculated into ten separate culture tubes containing 9.9 mL DM25 and incubated in an Innova 3100 water bath shaker (New Brunswick Scientific) overnight at 37.0° and 120 RPM. Cells from all ten tubes were pooled, and genomic DNA was extracted from these samples using Wizard Genomic DNA Purification Kits (Promega). This method of pooling independently derived cultures provided a way to later filter out any mutations that might have risen to high frequency during the process of recovery from being frozen. Our reasoning was that a specific mutation might arise in a single tube, but would be less likely to arise independently in multiple tubes, providing an empirical frequency cutoff of 10%, based on the number of independent tubes pooled. DNA from the Lazarus ancestor was extracted in the same manner, but pooled from four tubes rather than 10.

Genomic DNA libraries were prepared using the TruSeq DNA PCR-Free Library Preparation Kit (Illumina). The 26 Lazarus samples were multiplexed and sequenced in two lanes of an Illumina HiSeq 2500 in rapid mode at UC Irvine’s Genomics High-Throughput Facility. Two ancestral samples—one working stock (Figure 1) and one backup stock—were also sequenced on an Illumina HiSeq 3000 at the Bioinformatics Core Facility at the UC Davis Genome Center.

Mutations and mutation frequencies were called using breseq (Deatherage and Barrick, 2014) in polymorphism mode, using the *E. coli* B REL606 genome as a reference. Six regions (*topA, spoT, glmU/atpC, pykF,yeiB,* and the *rbs* operon) possess mutations that differ between REL606 and REL1206 (Barrick et al., 2009; Tenaillon et al., 2012), so there regions were excluded from our analyses. In theory, breseq provides information about duplications and deletions by reporting novel junctions. No evidence of novel junctions or sequencing coverage was found for large deletions in our data set, but breseq did provide some novel junction evidence for the presence of large duplications. To assess duplications more formally, we compared unique reads (mapping quality >5 in samtools 1.3) across 10 kb regions of the genome, defining duplications as regions with more than twice the average genome coverage.

**Fitness Assays:** Relative fitness values were assessed by performing standard competition assays (Lenski *et al.*, 1991; Tenaillon et al., 2012). Samples of frozen Lazarus populations and an ancestral stock containing a neutral Ara+ mutation (REL1207) were grown overnight in 9.9 mL DM25 using an Innova 3100 water bath shaker (New Brunswick Scientific) at 37.0° and 120 RPM. 100 μL of Lazarus culture was added to each of six replicate tubes containing 9.9 mL DM25. Likewise, 100 μL of ancestor culture was added to each of three tubes containing 9.9 mL DM25. These were incubated overnight at 42.2° and 120 RPM. All three ancestral tubes were pooled and vortexed. On day zero (t_0_) 75 μL of ancestor culture and 25 μL of a Lazarus culture were added to a tube of 9.9 mL DM25 and vortexed. 100 μL of this competition culture was diluted 1/100, and 100 μL of the dilution was plated on a TA plate for overnight incubation at 37.0°. The remaining competition culture was grown overnight in the water bath at 42.2° and 120 RPM. After one day of competition (t_1_), 100 μL of competition culture was diluted 1/10,000 and plated as above. Ara+ and Ara− colonies were counted for both t_0_ and t_1_. Fitness values were calculated using the methods of Lenski et al. (1991), but results were qualitatively identical using the methods of Chevin (2011).

**Data Availability:** All sequence data have been submitted to the NCBI short-read archive (Project Number XXXXXXX). Cell density measurements can be found in Table S1. Details about mutations present at >10% frequencies within a Lazarus population are reported in Table S2. Colony counts from fitness assays are reported in Table S3.

## RESULTS

**Frequency of Lazarus Events:** Over the course of five days, we measured the cell densities of 308 populations as they evolved (or more commonly, went extinct) at 43.0° (Table S1). 12 of these populations (one in one week, 11 in another week) were excluded from our final analysis due to mechanical issues during particle counting. Thus, 296 populations were considered for determining the frequency of Lazarus events. In total, we observed 26 Lazarus events (Figure 2), placing their frequency in our system at 8.8% (26/296), comparable to the frequency of 10% (3/30) observed by Mongold et al. (1999) in their preadapted populations grown at 44.0° (Mongold *et al.*, 1999). It is worth noting, however, that our populations were not pre-adapted to high temperature conditions. When we grew 88 populations for five days at 44.0°, none survived.

**Figure 2:**
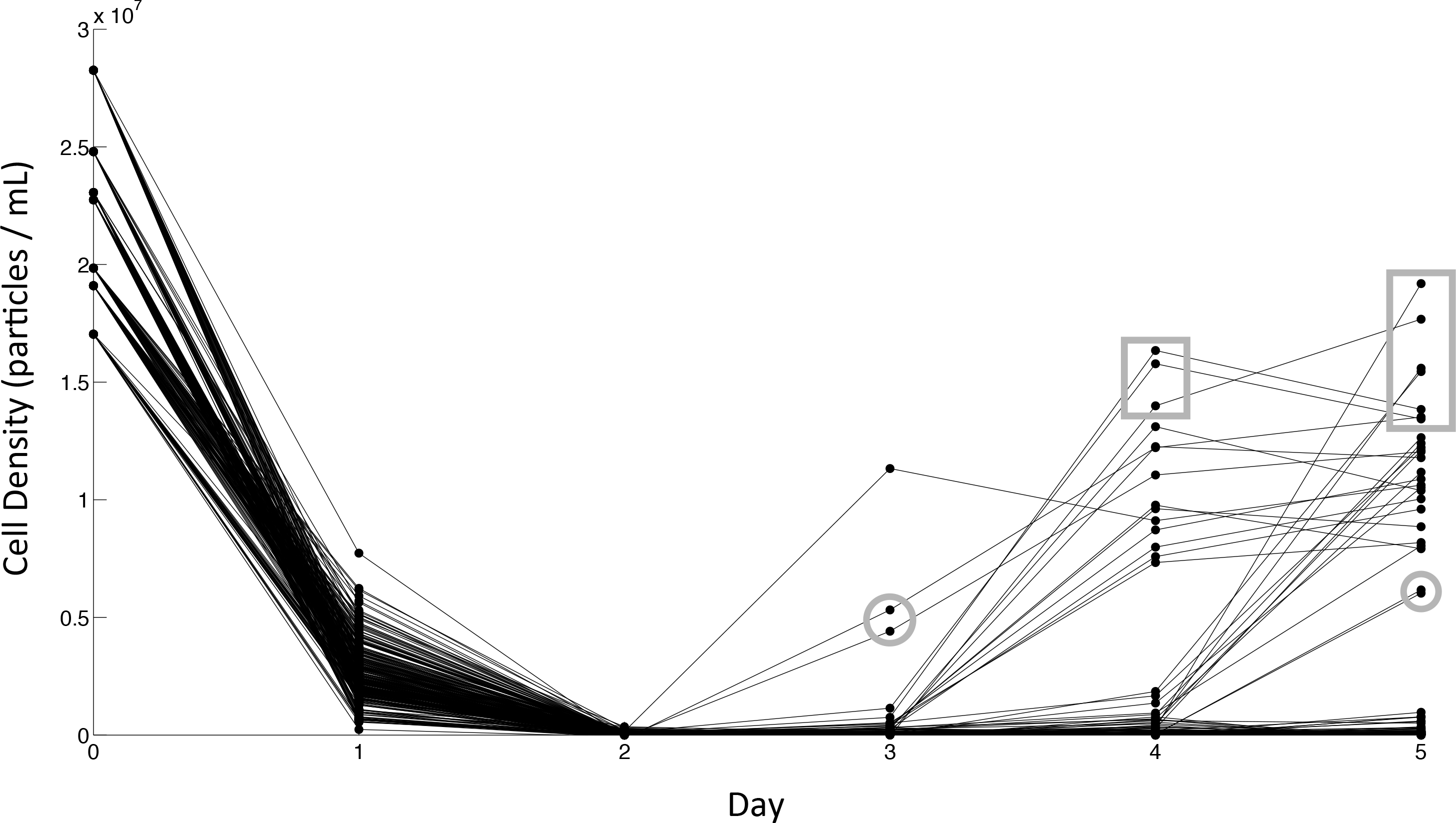
Population cell densities over time. Most populations went extinct over the course of five days. A total of 26 Lazarus events were observed across the third, fourth, and fifth days of growth. The timing of Lazarus events was determined by the day at which cell density increased by an order of magnitude over the previous day. Populations possessing *rpoBC* mutations are indicated by rectangles. Populations possessing duplications are circled.

Most Lazarus events occurred by day four, with events on day five also being common (Figure 2). Only three Lazarus events occurred on day three, suggesting a limit to how quickly beneficial mutations can become established within populations. Lazarus events occurred more often in some weeks than others. The highest incidence of Lazarus events in one week was 25% (11/44), and the lowest was 0% (0/33). Initial cell density bore no relationship to the number of Lazarus events observed in a given week (r = −0.24, *p* = 0.61), and initial cell density was not significantly correlated with the timing of the first population recovery in a given week (r = 0.50, *p* = 0.32).

**Mutations Associated with Lazarus Events:** To characterize the genomic changes underpinning Lazarus events, we sequenced each of the 26 Lazarus populations and identified the frequencies of mutations using breseq (Deatherage and Barrick, 2014). Among the 26 Lazarus populations, we identified 419 simple mutations (i.e., single nucleotide changes and small indels) within 122 unique genic and intergenic regions. Of these, 100 were called at frequencies >10% in their populations, and these comprised 32 unique genic and intergenic regions (Table 1, Table S2, Figure 3).

**Figure 3:**
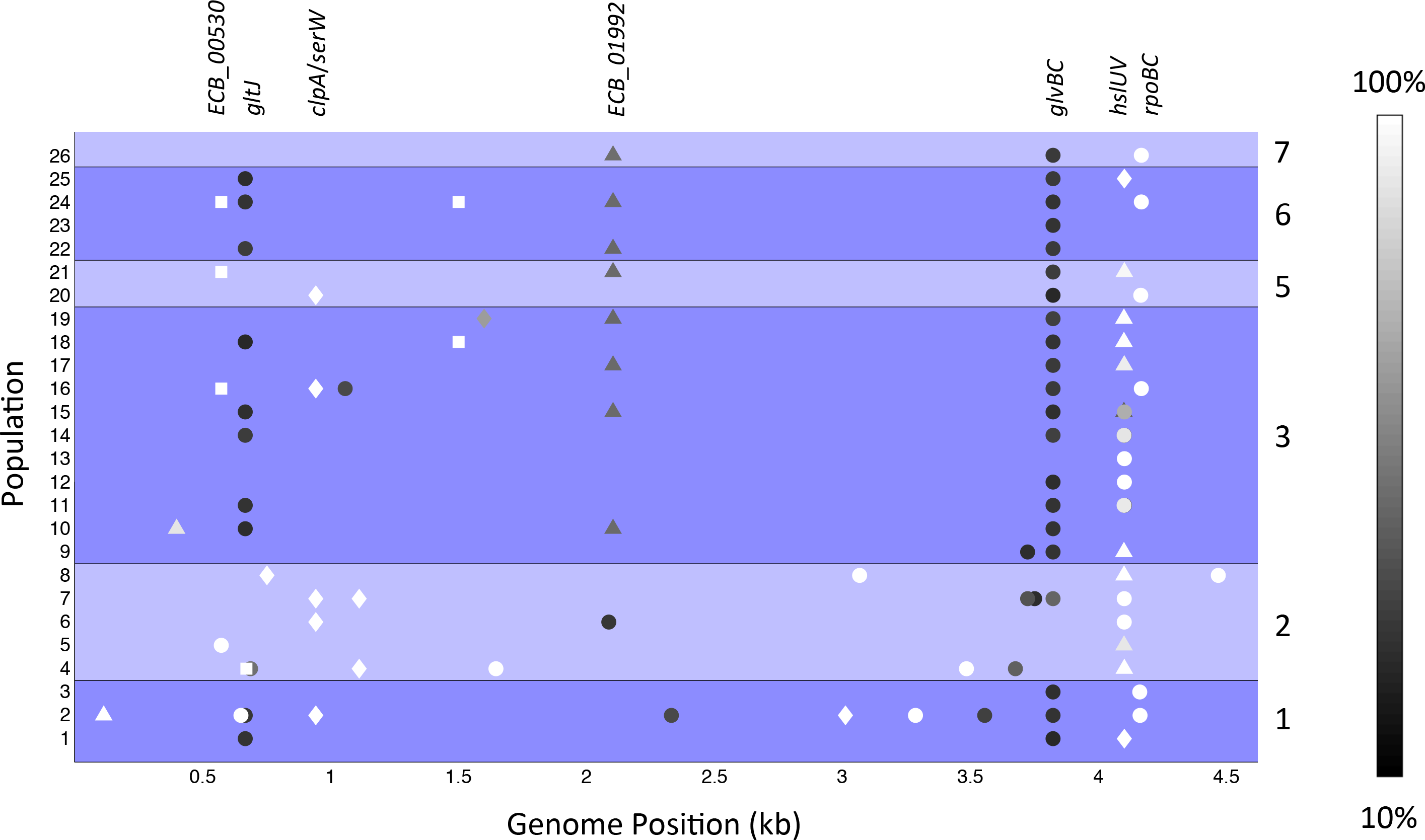
Genome-wide distribution of mutations in Lazarus populations. Different weeks are separated into groups and labeled at the right. Mutations are colored by their frequency in the population according to the scale at the right. Synonymous, nonsynonymous, indel, and intergenic mutations are represented by squares, circles, triangles, and diamonds, respectively. Only mutations at frequencies greater than 10% are shown. Mutations occurring in more than two populations are labeled at the top.

**Table 1:**
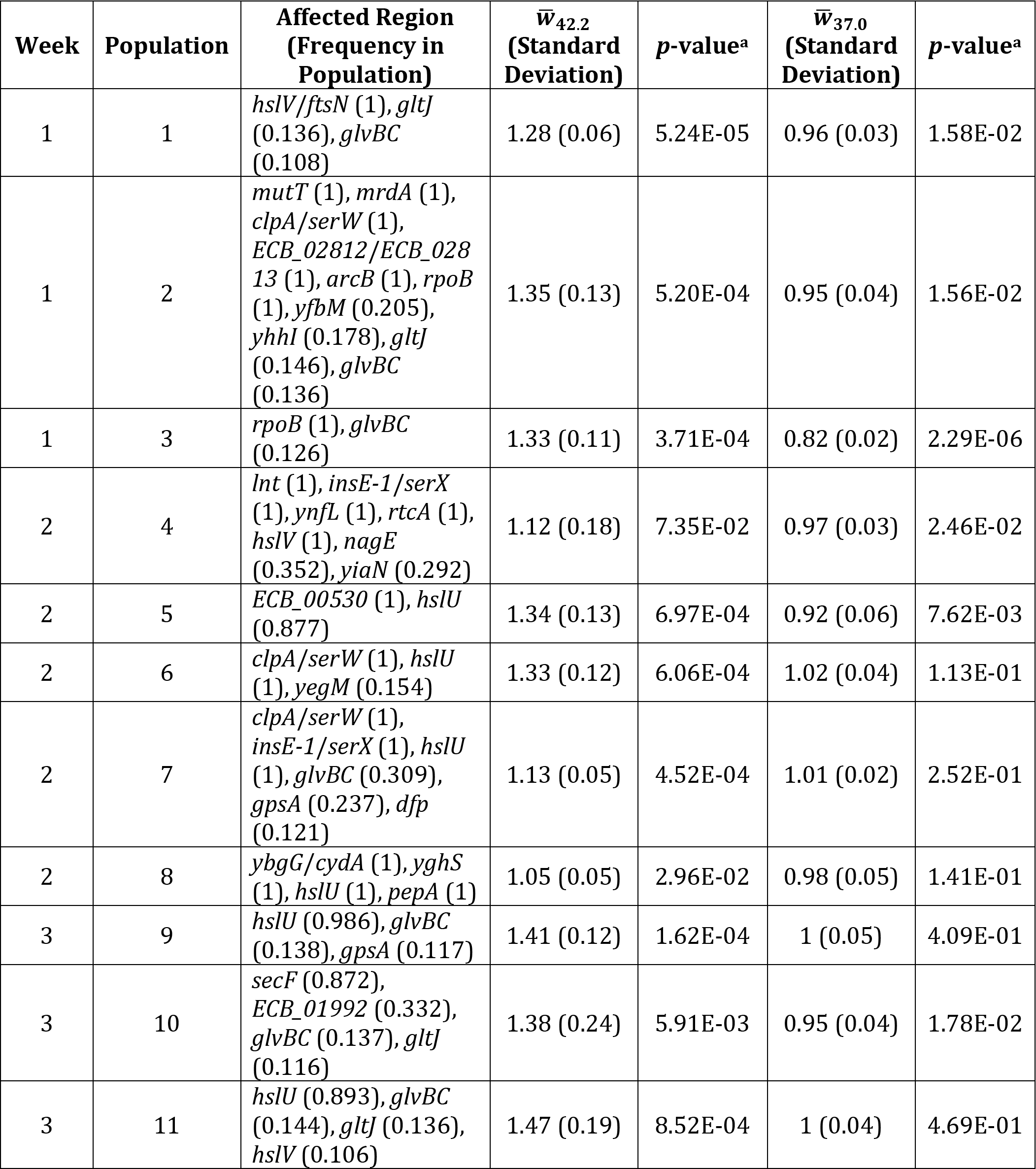
Mutations present in populations at frequencies >10% and mean fitness values (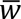) of populations relative to their ancestor at 42.2° and 37.0°.

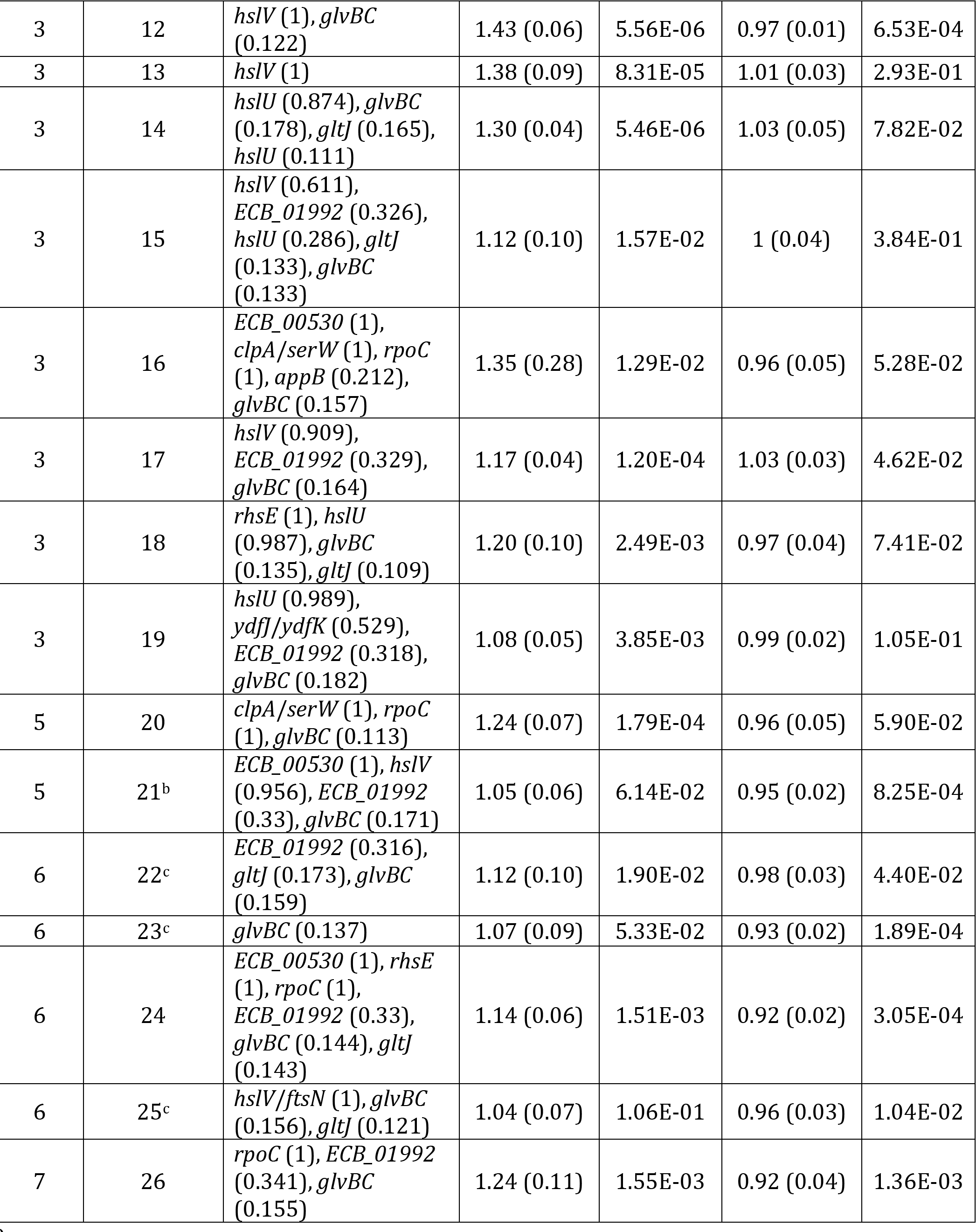

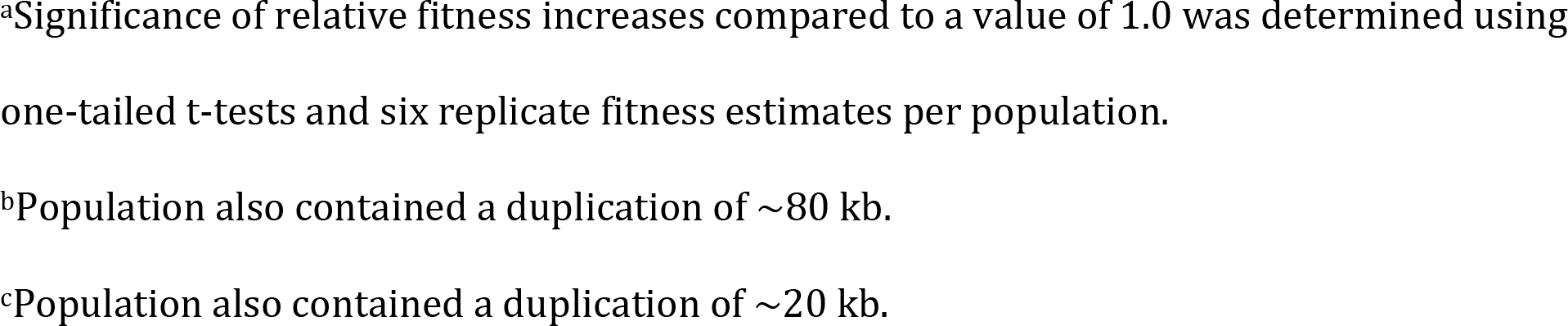

For a mutation to rescue a population from the brink of extinction, it must sweep through the population to an appreciable frequency. Focusing on mutations at frequencies > 85% in a population as “fixed”, we observed 46 fixed mutations within 20 unique genic and intergenic regions among 23 of the 26 Lazarus populations. On average, each population possessed 1.8 fixed mutations. 42% (11/26) of populations had just one fixed mutation, making these genetic changes the likely drivers of population recovery. Additionally, one of our Lazarus populations (#2; Table 1) evolved a mutator phenotype due to a small deletion in the *mutT* gene. Not surprisingly, this population contained six fixed mutations, the most of any population in the experiment.

Of the three remaining populations lacking fixed mutations, one (#15) possessed mutations in both *hslU* and *hslV* (see below) at intermediate frequencies, and two (#22 and #23) possessed large duplications at detectable frequencies. In total, four populations contained large duplications based on their sequencing coverage profiles, with three of these (#22, #23, and #25) occurring in parallel within a single week. These three parallel duplications included the genes *groEL* and *groES*, which encode a major chaperonin complex that is both necessary for growth under normal conditions and induced during growth at elevated temperatures (Fayet *et al.*, 1989; Hayer-Hartl *et al.*, 2016). The fourth population (#21) contained a duplication that includes some or all of the *hslUV* operon (see below).

To be assured that the fixed mutations arose as a consequence of thermal treatment, we also sequenced two control cultures. The first culture was inoculated from the frozen ancestor stock used throughout the experiment, and the second was inoculated from a reserve ancestor stock. We sequenced both cultures to >2,000x and identified no fixed (>85% frequency) mutations relative to the REL1206 genome. We did identify 1,105 potential variants across both cultures at an average frequency was ~1.3%. There were, however, three mutations that exceeded 10% frequency. Of these, one was shared with Lazarus populations: a four-nucleotide indel within the *ECB_01992* gene found at a frequency of 19% and 21% in the two ancestral samples. Interestingly, this mutation was found at frequencies between 32% and 34% across eight of the 26 Lazarus populations, but it never reached fixation. The lack of overlapping mutations between control and Lazarus populations (with the notable exception of the *ECB_01992* indel), combined with the low frequency of mutations in control populations, argues that the fixed mutations in Table 1 are indeed a consequence of population recovery under a temperature that is typically lethal.

**Genetic Targets of Parallel Mutations:** Experimental evolution derives its power from replication. If genetic changes occur multiple times across replicates, it provides strong evidence that these changes are adaptive (Woods *et al.*, 2006). In our experiment, we found consistent fixation within a pair of genes, *hslU* and *hslV*, which together comprise an operon that encodes a heat shock protease system (Missiakas *et al.*, 1996; Bochtler et al., 2000). Fixed mutations in *hslUV* affected 62% (16/26) of Lazarus populations: nine populations possessed fixed mutations in *hslU*, and another seven possessed fixed mutations in either *hslV* or its upstream region. We also noticed a striking pattern of parallelism in the *hslUV* operon, because parallel mutations occurred at the level of individual nucleotides within single weeks, but not between weeks (Figure 4A), and this parallelism was most prevalent in weeks that showed the most Lazarus events. These parallel events included small frameshift indels. In total, 69% (9/13) of *hslUV* mutations caused frameshifts, suggesting that interruption of *hslUV* function may be adaptive.

**Figure 4:**
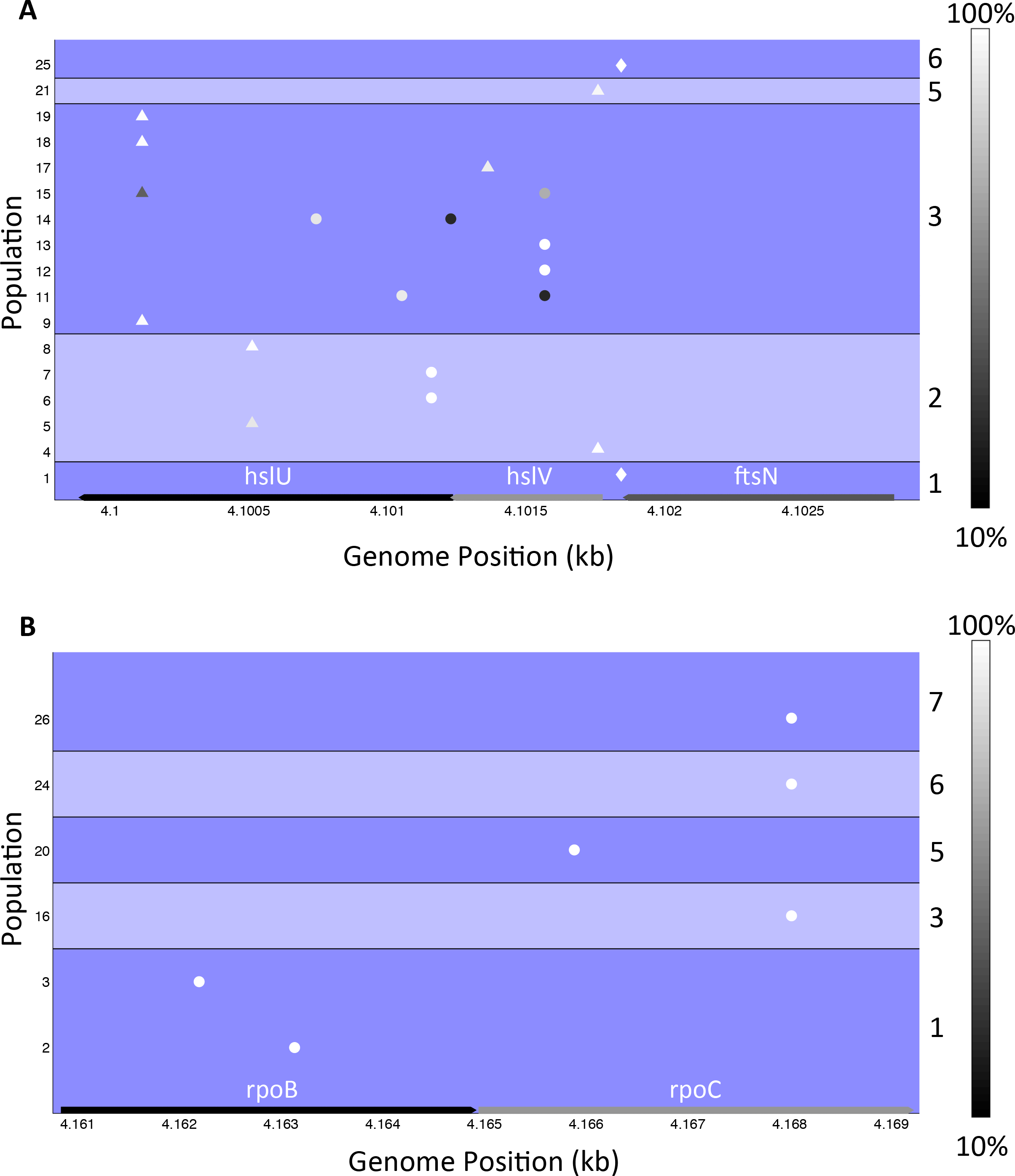
Distribution of mutations in the (A) *hslUV* and (B) *rpoBC* operons. Different weeks are separated into groups and labeled at the right. Mutations are colored by their frequency in the population according to the scale at the right. Only mutations at frequencies greater than 10% are shown. Synonymous, nonsynonymous, indel, and intergenic mutations are represented by squares, circles, triangles, and diamonds, respectively, as in Figure 3.

Like *hslUV*, the *rpoBC* operon also accumulated multiple fixed mutations (Figure 4B), which were found across six populations (#2, #3, #16, #20, #24, and #26). Four populations had fixed mutations in *rpoC*, another two had fixed mutations in *rpoB*, and all were nonsynonymous mutations. Interestingly, the populations bearing *rpoBC* mutations were distinct from those with *hslUV* mutations; in other words, no populations contained fixed mutations in both operons. Populations with fixed *rpoC* mutations also had nucleotide-level parallelism, but unlike *hslUV* mutations, this parallelism occurred across weeks, rather than within a single week (Figure 4B).

A third region, *clpA/serW*, accumulated the same fixed point mutation in five different populations (#2, #6, #7, #16, and #20) across four independent weeks. This region lies between *clpA*—one component of a protease system of similar function to that encoded by *hslUV* (Kwon *et al.*, 2004)—and *serW*, which encodes a serine-bearing tRNA.

Mutations within *hslUV, rpoBC*, and *clpA/serW* account for 59% (27/46) of fixed mutations. In many cases, *hslUV, rpoBC*, and *clpA/serW* mutations were fixed with other mutations. For example, population #4 had a total of five fixed mutations, including a mutation in *hslV.* Importantly, a single mutation in *hslUV* or *rpoBC* represents the *only* fixed mutation in several populations (#1, #3, #9, #11, #12, #13, #14, #17, #19, #25, and #26), suggesting that some mutations within these operons are sufficient to rescue populations and drive a Lazarus effect.

Finally, because Lazarus events occurred at different times during the course of a week, we were interested to see if there was any relationship between the timing of Lazarus events and the identities of fixed mutations, but we found no obvious relationship (Mann-Whitney U test comparing *hslUV* and *rpoBC* populations: *p* = 0.34).

**Density and Fitness:** The hallmark of a Lazarus event is the marked increase in cell density that occurs following a population crash, but not all populations rebound to the same final cell density (Table S1). We tested for a relationship between cell densities after population recovery and the fixed mutations present within those populations. At both days four and five, populations with fixed *rpoBC* mutations were among those with the highest cell densities. All six populations containing *rpoBC* mutations were among the seven populations with the highest cell densities at day five, and three more had the highest cell densities on day four. *rpoBC* populations had significantly higher cell densities than all other recovered populations on both days (Mann-Whitney U test: day four *p* = 0.0044, day five *p* = 3.71 × 10^−4^). In addition, populations with large duplications yielded cell densities approximately half as great as other populations that first experienced recoveries on the same day (Figure 2).

Given apparent relationships between mutations and cell densities, we quantified the relative fitnesses (*w*) of our Lazarus populations. Using standard competition assays (Lenski *et al.*, 1991; Tenaillon et al., 2012; Rodríguez-Verdugo *et al.*, 2014), we competed each Lazarus population against REL1207, a strain identical to our Lazarus ancestor (Table S3). These relative fitness assays were performed at 42.2° rather than 43.0°, because REL1207 populations are sustainable at the former temperature but crash at the latter, making *w* measurement impossible.

We estimated *w* each of the 26 populations. On average, the Lazarus populations showed a *w* increase of 24% relative to the ancestor, with 22 of 26 populations having *w* > 1.0 (*p* < 0.05, Table 1). Although the average *w* increase of the *rpoBC* populations (28%) exceeded that of all the other populations (22%) and the *hslUV* populations alone (23%), it was not significantly higher than either (one-tailed t-test, unequal variance: *p* = 0.15 and *p* = 0.18, respectively).

Strangely, although the relative fitness values of *rpoBC* populations were not statistically higher than those of *hslUV* populations, *rpoBC* populations produced significantly more colonies during the competition experiments than *hslUV* populations at both time points (t_0_ and t_1_) in the competition (one-tailed t-test, unequal variance: t_0_ *p* = 2.42 × 10^−5^, t_1_ *p* = 2.81 × 10^−4^). The ancestor colony counts were unaffected between these two groups (one-tailed t-test, unequal variance: t_0_ *p* = 0.42, t_1_ *p* = 0.49). While this did not translate into a difference in *w*, it did agree with the cell density measurements at day five.

We also assessed whether Lazarus populations showed any fitness tradeoffs at their ancestral growth temperature of 37.0×. On average, the Lazarus populations exhibited a fitness decrease of 3× relative to the ancestor, with 14 of 26 populations having relative fitness values significantly < 1.0 (one-tailed t-test: *p* < 0.05, Table 1). Populations with fixed *rpoBC* mutations had an average *w* decrease of 8% relative to the ancestor (one-tailed t-test: *p* = 0.0082), which was significantly lower than all other populations (one-tailed t-test, unequal variance: *p* = 0.019). Although some *hslUV* populations had fitness values significantly < 1.0 (Table 1), *hslUV* populations had an average fitness decrease of 1% relative to the ancestor, which was not significantly different from 1.0 (one-tailed t-test: *p* = 0.054), suggesting less of a fitness tradeoff for *hslUV* populations compared to *rpoBC* populations.

## DISCUSSION

We have performed experiments to characterize the genetic mutations that contribute to the phenomenon of rescue from otherwise lethal temperatures. Like Mongold et al. (1999), who previously studied the Lazarus effect, we find that rescue is infrequent, because it occurs in only 8.8% of experimental populations. It is nonetheless frequent enough that it has the potential to be a potent source of evolutionary innovation. Moreover, we have made a series of puzzling observations in the course of our experiment. The first is that there is a striking lack of independence among experiments, because we were more apt to find the same genetic mutations among populations within weeks, rather than between weeks. There were nonetheless notable parallels across weeks as well, suggesting that there is finite pool of potentially adaptive rescue mutations. The second is that there is a surprising amount of polymorphism within populations; even though there are clear ‘driver’ mutations that become fixed in Lazarus populations, we have detected additional polymorphic mutations at frequencies of >10% in most of our populations. Finally, there are clear patterns to the driver mutations, especially in relation to a previous thermal stress experiment at non-lethal temperatures (Tenaillon et al., 2012). We have identified multiple driver mutations in two sets of genes: those encoding subunits of RNA polymerase (*rpoB* and *rpoC*), and those encoding heat shock proteases (*hslU* and *hslV*). In the Discussion below, we consider these and other points further, culminating with a verbal model of the adaptive dynamics of Lazarus populations and potential mechanisms for population rescue.

**Adaptation to Lethal and Nonlethal Temperature:** One of the initial mysteries of the dynamics of population recovery was why populations adapted to high, but nonlethal temperatures did not concomitantly extend their upper thermal niche, thus enabling survival at even higher temperatures (Mongold *et al.*, 1999). Instead, lethal temperature conditions remained lethal for most populations, and recovery occurred primarily in populations already adapted to elevated temperature, suggesting that they were somehow pre-adapted to recover from high and otherwise lethal temperatures (Mongold *et al.*, 1999). Moreover, fitness measurements suggested that populations that recovered in lethal temperatures experienced tradeoffs at elevated, but nonlethal temperatures, implying that distinct sets of mutations are adaptive under nonlethal and lethal temperature regimes (Mongold *et al.*, 1999).

To determine if there is a distinction between the sets of mutations that arise under lethal and nonlethal conditions, we have compared the genetic data of our Lazarus experiments at 43.0° to the results of a previous long-term evolution experiment conducted at a sustainable 42.2° (Tenaillon et al., 2012). These experiments can be compared directly because they were based on the same ancestor (REL1206), grown in the same media (DM25), and under the same conditions (i.e., 10 mL of media, with 120 RPM shaking). Although there are other differences (see below), the major difference is between a high but sustainable temperature (42.2°) and a high but lethal temperature (43.0°).

Among the parallel results between experiments, one was the recovery of point mutations in *rpoB* and *rpoC*. The *rpoB* gene is a common target for mutations across a wide array of evolution experiments (Herring et al., 2006; Conrad et al., 2009; Charusanti et al., 2010), and it was also the most mutated gene in the 42.2° experiment (Tenaillon et al., 2012). In that experiment, 76 of 115 clones contained an *rpoB* mutation, another 21 of 115 clones contained an *rpoC* mutation, and a total of 5 of 115 lines contained a mutation in both genes. At least some of the *rpoB* mutations were likely to have been drivers of adaptation, both because they fixed rapidly during the course of the 42.2° experiment (Rodríguez-Verdugo *et al.*, 2013) and because they provided a ݾ22% fitness benefit, on average, at 42.2° as single mutations in the REL1206 background (Rodríguez-Verdugo *et al.*, 2014). Our Lazarus experiments also suggest that *rpoB* and *rpoC* mutations are adaptive, because we have detected two populations with fixed (>85% estimated frequency) *rpoB* mutations and four populations with fixed *rpoC* mutations (Table S2). Of these six mutations, we can definitively identify at least one *rpoB* and one *rpoC* as driving mutations, because they were the lone high frequency variants in the recovered population (#3 and #26; Table 1).

The two experiments also identify mutations in the *hslUV* operon, but the frequency of mutations differs. In the 42.2° experiment, only two of 115 (1.7%) clones contained a mutation in the *hslUV* region. Neither were obvious knockouts; one mutation was intergenic, and the other was a nonsynonymous replacement (I136N). In contrast, 16 of 26 (61%) of Lazarus populations contained a mutation in *hslUV* or their upstream region, several of which appear to be drivers because they are the lone fixed mutation in the population.

We can compare results between the two experiments more formally by considering the frequency of mutations within specific genes. For example, in the *hslU* gene, one mutation was observed out of 115 clones from the 42.2° experiment, for a frequency of 0.009 (1/115). Assuming this is the probability of an adaptive mutation occurring in this gene in the Lazarus experiment, we can test whether our Lazarus observation of nine mutations events in 26 populations is expected. We find that the probability of observing nine or more *hslU* mutations in 26 observations is vanishingly small given a frequency of 0.009 (binomial: *p* = 7.77 × 10^−13^; Table 2). We note, however that this result changes somewhat if one assumes that the 26 populations are not independent (see below), and instead rely on the fact that *hslU* mutations occur in two of seven independent weekly trials (binomial: *p* = 0.02; not significant after Bonferroni correction; Table 2).

**Table 2:**
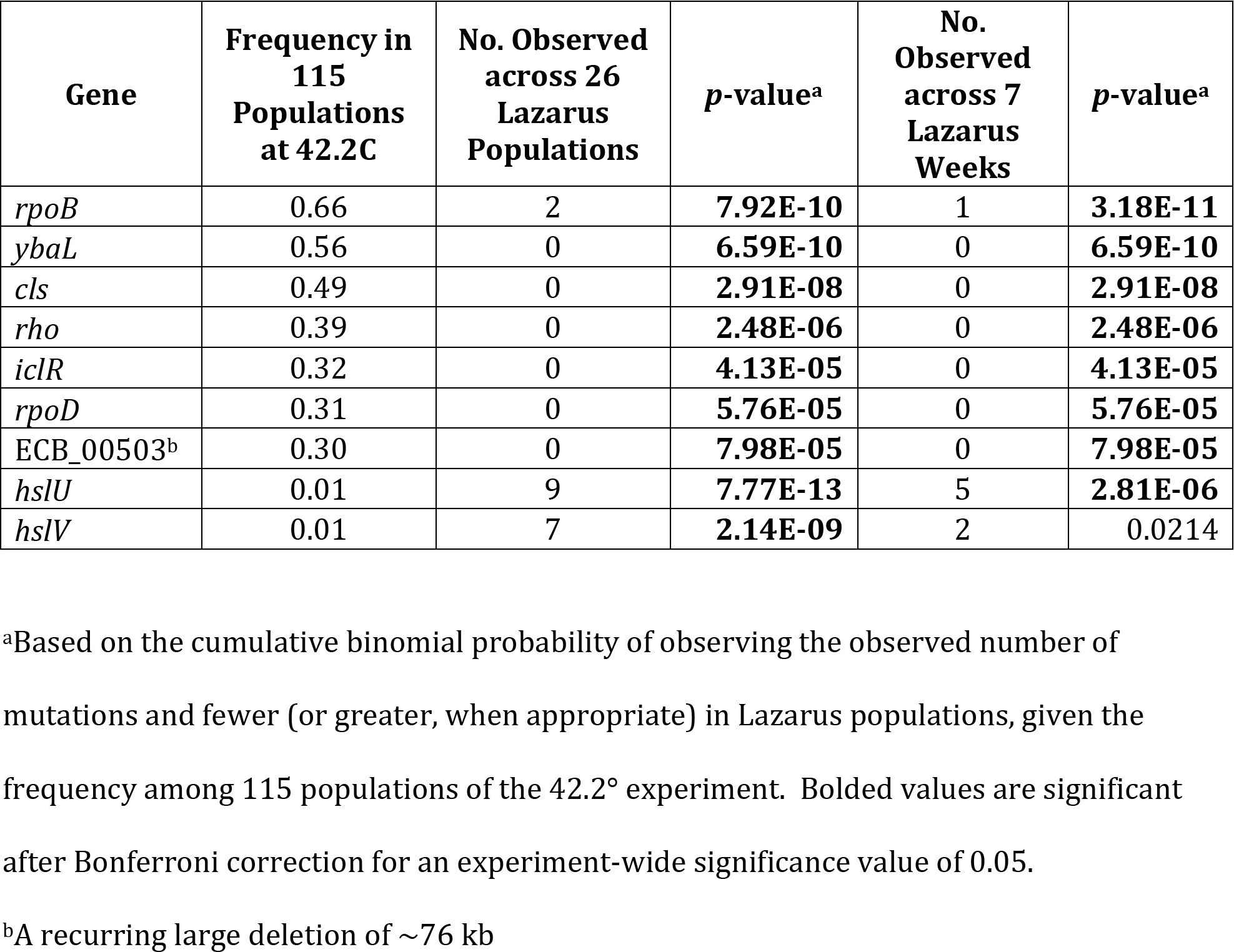
Mutations within specific genes that differ significantly in frequency between lethal and non-lethal high-temperature experiments.

We find that many common mutations differ significantly between the two experiments (Table 2), most because they have occurred too infrequently in the Lazarus populations (e.g., *rpoB,ybaL, cls, rho, iclR*, and *rpoD* mutations), but others because they occur too often in Lazarus populations (e.g., *hslV*). Notably, *rpoB* and *ybaL*, the two most mutated genes within the 42.2° experiment—and the *rho* and *cls* genes, which together define a second path of adaptation to 42.2° adaptation—are significantly underrepresented in Lazarus populations. One important exception to this trend is *rpoC*, because we cannot reject the hypothesis that *rpoC* mutations have occurred with equal frequency in the two experiments (binomial: *p* = 0.47 for 26 populations, *p* = 0.24 for 7 weeks). There are caveats to using this binomial approach for comparing experiments: specifically, some mutations are not strictly independent because they occur together on haplotypes, and the test relies on a single point estimate of probability from the 42.2° experiment. Nonetheless, the overarching impression is that the two experiments identify different sets of genes as loci for adaptive mutations, with the notable exception of *rpoC*.

Why might the sets of mutations differ between these two experiments? As we have noted, one difference between experiments is a modest but physiologically critical difference in temperature, but they also differ in at least three other features. The first is the method in which the experiments were started—single clones for each of the 115 populations at 42.2° (Tenaillon et al., 2012; Hug and Gaut, 2015) but a batch culture for each week of the Lazarus experiment. These batch cultures likely introduced nonindependence among replicates within weeks, but this non-independence is helpful for ultimately understanding the dynamics of rescue (see below).

A second difference is population size. Because REL1206 growth is sustained at 42.2°, populations remained large (~10^6^ individuals) for all phases of that experiment, whereas Lazarus populations initially crashed to undetectable or nearly undetectable levels (Figure 2). Within crashed populations, genetic drift may rapidly elevate the frequency of rare advantageous mutations, ultimately leading to an acceleration of the rate of adaptation (Lang et al., 2013). We revisit this point further below.

Finally, the two experiments differ in duration; the 42.2° experiment ran for a year, but the Lazarus populations were propagated for only five days. This time difference is likely important, because large-effect mutations tend to be fixed early in experimental populations (Kryazhimskiy *et al.*, 2014). After the initial fixation of large-effect mutations, the adaptive fixations that follow tend to have incrementally diminishing fitness benefits (Chou *et al.*, 2011; Khan *et al.*, 2011; Kryazhimskiy *et al.*, 2014). Given the short timespan of Lazarus experiments, we are likely uncovering only large-effect mutations.

Unfortunately, we cannot compare them directly to early large-effect mutations in the 42.2° experiment, because it is not yet known which mutations were fixed early in the that experiment (with the notable exception of *rpoB* mutations) (Rodríguez-Verdugo *et al.*, 2013). If we were to continue the Lazarus experiment, it is possible that the set of adaptive mutations separating lethal and nonlethal temperatures would diminish (Table 2). It is important to note, however, that time alone does not explain differences between the experiments, because mutations within *hslV* are common, fixed drivers in several Lazarus populations but were rare in the year-long 42.2° experiment.

***hslUV, rpoBC,* and the Roles of Standing Variation and Genetic Drift:** *hslUV* and *rpoBC* mutations drive adaptation and population recovery, but the evolutionary dynamics of mutations in these operons differ substantially for three reasons. First, *hslUV* and *rpoBC* mutations are not found together as fixed in the same population, which is statistically improbable given their respective frequencies across populations and given that at least one is fixed as a putative driver in each population with two or more fixed mutations (*p* < 0.02). The lack of co-occurring *rpoBC* and *hslUV* mutations suggests an initial canalization of the adaptive response, focused either on a pathway through heat shock proteins or through RNA polymerase, but not through both. Such a pattern is consistent with negative epistasis observed between the *rho* and *rpoB* pathways of the 42.2° experiment (Tenaillon et al., 2012). These results imply that epistasis shapes trajectories even at the earliest stages of the adaptive process. Second, as noted above, their patterns of parallelism differ. Specific *hslUV* mutations tend to repeat across populations within weeks, but *rpoBC* mutations repeat across weeks (Figure 4)—for example, the same nonsynonymous mutation in *rpoC* (W1020G) occurred in weeks three, six, and seven. Finally, populations with *rpoBC* vs. *hslUV* mutations had distinct fitness patterns. Based on cell densities and relative fitnesses at 42.2° (Table 1), mutations in the *rpoBC* operon offer an advantage that is at least equivalent to, if not greater than, mutations in the *hslUV* operon. Despite this, *hslUV* mutations were more prevalent across all populations.

Together these observations suggest different evolutionary dynamics for mutations within *hslUV* and *rpoBC*. One possibility is that these two adaptive strategies are associated with different costs under non-stressful conditions. To assess this possibility, we competed Lazarus populations against the REL1207 ancestor at 37.0°. Our results indicate that the *hslUV* populations do not differ statistically as a group in their fitness from the ancestor, suggesting that *hslUV* mutations are neutral (or only mildly deleterious) at 37.0°. This observation may not be surprising, considering that the inactivation of heat shock genes is unlikely to be critical at 37.0°. In contrast, *rpoBC* populations have a greater fitness tradeoff (significantly lower fitness) at 37.0° than *hslUV* populations, suggesting that *rpoBC* mutations are deleterious at 37.0°.

Together, these lines of evidence suggest a model for the establishment of recovery. This model relies critically on the fact that the replicates within a week were established from batch cultures at 37.0° (Figure 1). Previous work in nonlethal experimental evolution systems has suggested that adaptation is not mutation-limited (Lang *et al.*, 2011), and we believe that to be the case here. Starting with our final cell densities from day zero (acclimation growth in DM25 at 37.0°), using the expected 6.64 generations produced in DM25 (Lenski *et al.*, 1991), and assuming an *E. coli* mutation rate of 10^−3^ per genome per generation (Lee *et al.*, 2012), we can calculate the number of mutations expected at each nucleotide position during DM25 acclimation. Using a conservative six generations of binary fission, our populations would have sampled between 0.36 and 0.60 mutations per nucleotide, with an average of 0.47. It is important to note that these are only the mutations that could have arisen during growth in DM25; an additional eight hours of growth took place in LB prior to this (Figure 1).

Based on the length of the *hslUV* and *rpoBC* operons, we expect that mutations in *rpoBC* occur more often than in *hslUV*. However, several effects may have acted individually or together to generate a bias toward *hslUV* mutations. First, many *hslUV* mutations constituted frameshifts, suggesting that a simple loss of function of this operon was beneficial. Therefore, while *hslUV* might have been less likely to be mutated than *rpoBC* based on length, more mutations may result in a fitness advantage. In contrast, *rpoBC* mutations must retain function, suggesting there may be fewer potentially adaptive mutations from which to sample. Second, as noted above, populations with mutations in *hslUV* did not exhibit antagonistic pleiotropy to the same degree as populations with *rpoBC* mutations; they did not have an associated fitness cost at the acclimation temperature of 37.0°, suggesting *hslUV* mutations are more likely to exist in batch culture and may reach frequencies that enhance the possibility of sampling into separate populations, perhaps explaining the high degree of parallelism across populations within weeks. By contrast, because individuals with *rpoBC* mutations are less fit than the ancestor at 37.0°, new *rpoBC* mutations in batch culture would tend to be lost rapidly, making them less likely to be sampled and distributed to individual Lazarus tubes. This argument presumes that much of the initial selection to high temperature acts on standing variation that was generated at 37.0°.

If selection acts on standing variation and *rpoBC* mutations are deleterious at 37.0°, how do any *rpoBC* mutations survive? There are two possibilities. The first is that the *rpoBC* mutations occur *de novo* after transfer to lethal temperature. Under this hypothesis, such mutations would be expected to be rare, because the population size decreases immediately after introduction to high temperature. If this is true, however, one might predict that *rpoBC* populations recover later than other populations, and this is not the case. The second possibility is that the population crash increases genetic drift, occasionally establishing a rare, highly advantageous allele for subsequent fixation.

**Mechanism:** We have shown that population recovery occurs rarely and that it tends to be accompanied by mutations in *hslUV* or *rpoBC*, but not both. We believe that the pattern of parallelism within and between weeks is driven in part by the antagonistic pleiotropy of *rpoB* mutations and the relative lack thereof for *hslUV* mutations. But we have failed to address another important question: what is the molecular mechanism by which these mutations permit population recovery?

There are already some insights into the potential benefits of *rpoB* mutations at high temperature, because similar *rpoB* mutations have been shown to have large effects on gene expression at high temperature (Rodríguez-Verdugo et al., 2016). For example, single mutants with *rpoB* affect the expression of thousands of genes at high temperature, and these shifts tend to move gene expression from a stress pattern to a state more like that of the ancestor (Rodríguez-Verdugo *et al.*, 2016). Accompanying these wholesale shifts in gene expression is a trend toward increasing transcriptional efficiency, suggesting that at least some adaptive *rpoB* mutations slow the polymerase and enhance termination at high temperature (Rodríguez-Verdugo *et al.*, 2016). We have no evidence, however, that the distinct Lazarus mutations in *rpoB* and *rpoC* confer these same benefits.

How mutations in *hslUV* enable population recovery is more of a mystery. Given that most mutations in the operon are frameshifts, they likely trigger a loss of function of the heat shock protease system. In the short term, this poses a problem, because these genes are normally upregulated upon the onset of heat shock conditions, although only weakly (Chuang *et al.*, 1993). Over a longer period of thermal stress, they are downregulated to below pre-stress levels (Rodríguez-Verdugo *et al.*, 2016). If *hslUV* plays a roll in the initial heat shock response and is downregulated later, the benefit from knock out mutations is perplexing. Perhaps the function of the hslUV-encoded protease system— to degrade both misfolded and properly folded proteins (Miller *et al.*, 2013)—becomes detrimental to the cell under extreme stress. Three lines of evidence support this idea. First, the *hslUV* protease system is known to target σ^32^, the major heat shock sigma factor of the cell, and therefore the absence of the *hslUV* proteases may enhance the production of other heat shock proteins (Kanemori *et al.*, 1997). Second, we observed three mutants (#22, #23, and #25) with large duplications centered on the *groEL* and *groES* genes, which encode a chaperonin complex important to the heat shock response (Richter *et al.*, 2010), and which are among the heat shock proteins upregulated in *hslUV* deletion mutants (Kanemori *et al.*, 1997). Lastly, our third-most common fixed mutation impacted the downstream region of *clpA*, a subunit of the *clpAP* protease system that has similar functions to, and overlapping substrate specificities with, *hslUV*(Kwon *et al.*, 2004). It is conceivable that altering the downstream region of *clpA* could result in its downregulation or a change in its function, which would produce similar outcomes to knocking out *hslUV*. Ultimately, we cannot yet ascribe a mechanism, but our hypothesis—i.e., that *hslUV* knockouts may indirectly lead to upregulation of other heat shock proteins through an effect on σ^32^—merits further attention.

## ACKNOWLEDGEMENTS

We thank R. Gaut and P. McDonald for contributing to data generation. S. Hug was supported by the National Institute of Biomedical Imaging and Bioengineering, National Research Service Award EB009418 from the University of California, Irvine, Center for Complex Biological Systems. The work was supported by National Science Foundation grant DEB-0748903.

